# Metabolic potential and survival strategies of microbial communities across extreme temperature gradients on Deception Island volcano, Antarctica

**DOI:** 10.1101/2020.08.07.241539

**Authors:** Amanda Gonçalves Bendia, Leandro Nascimento Lemos, Lucas William Mendes, Camila Negrão Signori, Brendan J. M. Bohannan, Vivian Helena Pellizari

**Affiliations:** Departamento de Oceanografia Biológica, Instituto Oceanográfico, Universidade de São Paulo (USP). Praça do Oceanográfico, 191. CEP: 05508-900 São Paulo, SP, Brazil; Laboratório de Biologia Celular e Molecular, Centro de Energia Nuclear na Agricultura, Universidade de São Paulo. Avenida Centenário 303, CEP 13416-00 Piracicaba, SP, Brazil; Department of Biology, Institute of Ecology and Evolution, University of Oregon, Eugene, OR, United States

**Author notes:** Corresponding author: Amanda Gonçalves Bendia, Address: Departamento de Oceanografia Biológica, Instituto Oceanográfico, Universidade de São Paulo, São Paulo, Brazil. Praça do Oceanográfico, 191. CEP: 05508-900 São Paulo, SP, Brazil.

**Keywords:** Deception Island, Antarctica, Microbial ecology, Microbial processes, Metagenome-assembled genomes, Extremophiles

## Abstract

Active volcanoes in Antarctica, in contrast to the rest of the icy landscape, have remarkable temperature and geochemical gradients that could select for a wide variety of microbial adaptive mechanisms and metabolic pathways. Deception Island is a stratovolcano flooded by the sea, resulting in contrasting ecosystems such as permanent glaciers (<0 °C) and active fumaroles (up to 100 °C). Steep gradients in temperature, salinity and geochemistry over very short distances have been reported for Deception Island, and have been shown to effect microbial community structure and diversity. However, little is known regarding how these gradients affect ecosystem functioning, for example due to inhibition of key metabolic enzymes or pathways. In this study, we used shotgun metagenomics and metagenome-assembled genomes to explore how microbial functional diversity is shaped by extreme geochemical, salinity and temperature gradients in fumarole and glacier sediments. We observed that microbial communities from a 98 °C fumarole harbor specific hyperthermophilic molecular strategies, as well as reductive and autotrophic pathways, while those from <80 °C fumaroles possess more diverse metabolic and survival strategies capable of responding to fluctuating redox and temperature conditions. In contrast, glacier communities showed less diverse metabolic potentials, comprising mainly heterotrophic and carbon pathways. Through the reconstruction of genomes, we were able to clarify putative novel lifestyles of underrepresented taxonomic groups, especially those related to Nanoarchaeota and thermophilic ammonia-oxidizing archaeal lineages. Our results enhance understanding of the metabolic and survival capabilities of different extremophilic lineages of Bacteria and Archaea.

## Introduction

The study of life in extreme environments has long fascinated biologists. Understanding how life persists at environmental extremes provides insight into how living systems function, as well as providing a unique window into the evolutionary history of life itself (Merino et al., 2019). The Deception Island volcano contains a unique combination of extreme temperatures and geochemical energy sources that together have the potential for selecting a wide variety of microbial adaptive mechanisms and metabolic pathways. Deception Island is located in the South Shetland Islands at the spreading center of the Bransfield Strait marginal basin, which harbors contrasting ecosystems of permanent glaciers and active fumaroles with continuous emissions of gases, mostly carbon dioxide and hydrogen sulfide (Somoza et al., 2004). This combination of glaciers and fumaroles is produced by the interaction between the cryosphere and water mass contact with hot ascending magmas (Geyer et al., 2019). Unlike Antarctic continental volcanoes, Deception Island fumaroles reach up to 100 °C and have direct marine influence, creating a remarkable combination of thermal, geochemical and salinity gradients (Bartolini et al., 2014; Herbold et al., 2014; Muñoz-Martín et al., 2005).

While early research carried out on Deception focused primarily on obtaining bacterial isolates from hot or cold ecosystems (e.g. Carrión et al., 2011; Llarch et al., 1997; Stanley et al., 1967), a more recent study was able to recover both psychrophilic and thermophilic isolates among the steep temperature gradients (Bendia et al., 2018a). Previous molecular studies described microbial diversity on Deception fumaroles using DGGE (Amenábar et al., 2013; Muñoz et al., 2011) and shotgun metagenomics to characterize the resistome profiles in cold sediments from Whalers Bay (Centurion et al., 2019). These previous studies were limited with respect to sampling depth and extent since only fumaroles or cold sediments were analyzed.

Two previous studies have focused on understanding the effect of Deception temperature gradients on microbial communities. The first, performed by our group, focused on determining taxonomic diversity through 16S rRNA gene sequencing (Bendia et al., 2018b), and a second study applied the Life Detector Chip (LDChip) to describe general functions of communities from Cerro Caliente (Lezcano et al., 2019). Our previous study showed that the steep gradients on Deception were able to select a unique combination of taxonomic groups found in deep and shallow hydrothermal vents (including hyperthermophilic Archaea, such as *Pyrodictium spp.*), geothermal systems and those typical from polar ecosystems (Bendia et al., 2018b). Also we reported that the bacterial community structure on Deception Island is strongly niche driven by a variety of environmental parameters (temperature, pH, salinity and volcanic geochemicals, such as sulfate), while archaeal diversity is mainly shaped by temperature (Bendia et al., 2018b). These previous studies, however, did not address the linkages between microbial structure and their adaptive and metabolic strategies over the particular environmental gradients found on Deception Island.

Although several studies have demonstrated that temperature is a primary driver of microbial taxonomic diversity in different geothermal and hydrothermal ecosystems (e.g. Antranikian et al., 2017; Price and Giovannelli, 2017; Sharp et al., 2014; Ward et al., 2017), it is still unclear to what extent temperature affects the functional processes of a microbial community, such as their adaptive mechanisms and metabolic pathways. The majority of these previous studies have focused on nonpolar thermal ecosystems, where the temperature range is narrower than in polar volcanoes; the exception is a study of deep-sea hydrothermal vents, in which the contrasting temperatures are created by the contact of heat with the surrounding cold seawater (with temperatures 0-4 °C). Indeed, the Deception communities from fumaroles have similar members (Bendia et al., 2018b) to those found in deep-sea hydrothermal vents (e.g. Dick, 2019; Nakagawa et al., 2006; Takai et al., 2001), which suggests that, regarding differences in pressure (and other environmental effects), the wide temperature range typical of polar volcanoes and deep-sea hydrothermal vents can act as a strong selective pressure that favors (hyper)thermophilic specialists capable of thriving in high temperatures but that can also tolerate the cold surroundings.

Furthermore, there is controversy about the impact of extreme environments on microbe-microbe interactions. Although some studies have reported a decrease in the frequency of microbe-microbe interactions inferred from co-occurrence patterns (Cole et al., 2013; Merino et al., 2019; Sharp et al., 2014), others have observed the opposite trend (Lin et al., 2016; Mandakovic et al., 2018). The analysis of co-occurrence patterns is useful for examining the nature of the ecological rearrangements that take place in a microbial community facing contrasting environments (Freilich et al., 2010; Mandakovic et al., 2018).

In the current study, we assessed the microbial functional profile in fumarole and glacier sediments from two geothermal sites on the Deception Island volcano, Antarctica. For this, we performed shotgun metagenomics to unveil functional diversity and reconstructed genomes to reveal the microbes` putative lifestyles and survival capabilities, combining the genomic information with community functional profiles. Here, we hypothesize that (i) similar to what has been reported for community diversity, survival and metabolic strategies are also influenced by the combination of geochemical, salinity and extreme temperature gradients; (ii) these communities follow the redundancy of metabolic potential that was reported for deep-sea hydrothermal vent communities; and (iii) microbe-microbe interactions decrease with increasing temperature. This study adds important information regarding the ecological processes of microbial communities inhabiting a steep gradient of temperature, and addresses central questions regarding the functional adaptability of extremophiles in polar regions.

## Materials and methods

### Study site and sampling strategy

Deception Island (62°58′ S, 60°39′ W) is a complex stratovolcano located in the South Shetland Islands, Bransfield Strait, near the Antarctic Peninsula. A past eruption occurring approximately 10,000 years ago collapsed the central part of the island giving rise to a flooded caldera called Port Foster Bay, 9 km in diameter (Baker et al., 1975). Fumaroles are found mainly at Fumarole Bay (FB), Whalers Bay (WB), and Pendulum Cove (Fermani et al., 2007; Geyer et al., 2019), and they are distributed mostly in submerged and partially submerged regions (intertidal zones), with temperatures varying from 40-60°C in WB and 80-100°C in FB (Rey et al., 1995; Somoza et al., 2004). Carbon dioxide and hydrogen sulfide gases are emitted by fumaroles and are oxidized to products such as sulfite and sulfate (Somoza et al., 2004; Zhang and Millero, 1993).

Sampling was performed during the XXXII Brazilian Antarctic Expedition (December 2013 to January 2014), with logistical support from the polar vessel NPo. Almirante Maximiano. We collected surface sediment samples (*ca.* 5 cm) in fumaroles and glaciers at geothermally active sites in FB (62°58’02.7” S, 60°42’ 36.4” W) and WB (62°58’45.1” S, 60°33’27.3” W) (Figure 1a and 1b), with temperatures between 0 and 98 °C. At each site, we obtained samples from three different points within the temperature gradient, and triplicates were performed for each collected point, totaling 18 sediment samples. Points A and B were defined as samples collected in fumaroles, while point C samples were collected from the glacier, few cm below the glacier’s edge (Figure 1c and 1d). The point FBA was the hottest fumarole, measuring 98 °C at the sediment surface (*ca.* 20 cm). Distances between fumaroles and glaciers at each site were approximately 15 m, and the WB and FB transects were approximately 10 km apart. All fumaroles were in the intertidal zone, except for point B from FB, which was in the subtidal zone (submerged at 50 cm depth in the water column). Samples were stored at −20 °C until arrival at the University of São Paulo, Brazil in April 2014.

**Figure 1.**
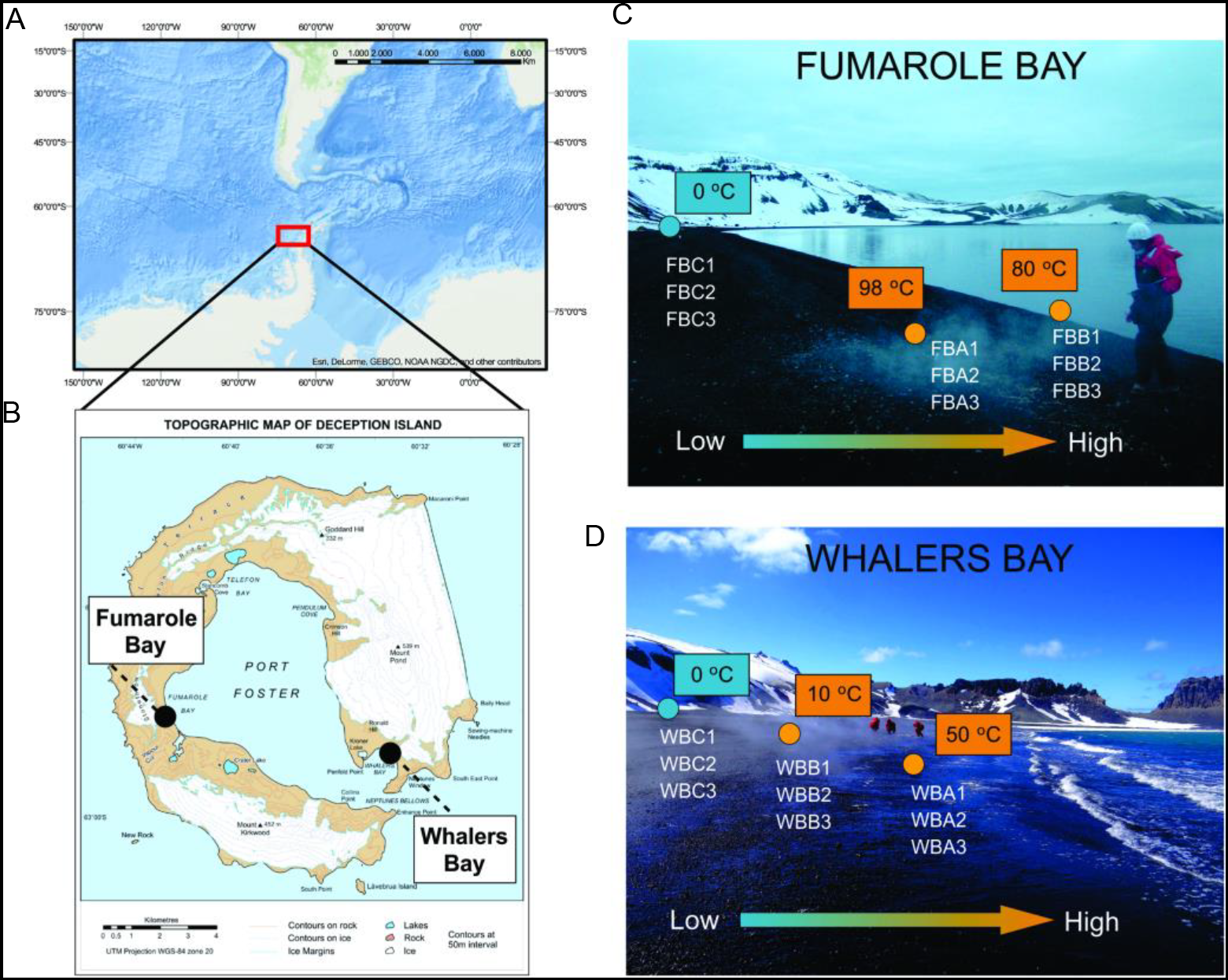
The sampling map with the location of Antarctic Peninsula (A) and Deception Island, with Fumarole Bay and Whalers Bay geothermal sites highlighted (B). Distribution of collected samples across environmental gradients at studied geothermal sites are described in C for Fumarole Bay and D for Whalers Bay. *In situ* temperatures are represented in blue (glaciers) and orange (fumaroles). The arrow indicates the direction of low and high values of temperature, salinity and volcanic compounds, such as sulfate. Figure was retrieved from Bendia et al., 2018b.

### DNA Extraction and metagenomic sequencing

To compare our results with the study performed by Bendia et al. (2018b), we used the same dataset and DNA extractions as a template for the metagenomics. Due to low DNA mass even after several concentration efforts, extracted DNA was subjected to multiple displacement amplification (MDA) using the illustra GenomiPhi V2 DNA amplification kit (GE Healthcare, Piscataway, NJ, USA), following the manufacturer’s instructions. Three amplification reactions per sample were pooled to obtain sufficient DNA for sequencing. DNA was then purified using AMPure XP beads kit (Beckman Coulter) following the manufacturer’s instructions. Library constructions and shotgun metagenomic sequencing were conducted at “Laboratório Central de Tecnologias de Alto Desempenho em Ciências da Vida” (LaCTAD), Universidade Estadual de Campinas State (UNICAMP), on the Illumina Hiseq 2000 platform using 2×100 bp paired-end system.

### Physical-chemical parameters

To correlate biological and environmental data, we used the physical-chemical parameters for sediments measured by Bendia et al. (2018b), which included granulometry, electrical conductivity, humidity, micronutrients (B, Cu, Fe, Mn, and Zn), organic matter, organic carbon, pH, P, Si, Na, K, Ca, Mg, Al, total nitrogen, nitrate, ammonia, and sulfate.

### Taxonomic and functional inference of metagenomic reads

Reads were quality trimmed using Sickle (Joshi and Fass, 2011) with phred >30 and then uploaded to MG-RAST (Keegan et al., 2016). Functional and taxonomic profiles of reads were generated through subsystem and best hit classifications using the SEED subsystem, M5NR (non-redundant protein database) and KEGG, available in MG-RAST (Aziz et al., 2008; Kanehisa and Goto, 2000; Keegan et al., 2016; Wilke et al., 2012), with the following parameters: 1×10^−5^ e-value, minimum 50 bp alignment, and 60% identity. Data generated by MG-RAST were statistically analyzed using Statistical Analysis of Metagenomic Profiles (STAMP) software (Parks et al., 2014) and R software (R Development Core Team), using the packages *vegan* (Oksanen, 2007) and *ggplot* (Wickham, 2011). The *p* values were calculated using Fisher’s exact two-sided test and the confidence intervals were calculated using the method of Newcombe-Wilson. Statistical comparisons were performed by grouping the samples according to environmental temperatures: glaciers, fumaroles up to 80 °C and fumarole at 98 °C. Principal component analysis (PCA) ordination was performed by using level 3 functions of SEED subsystems and then visualized in STAMP software. Values were normalized to relative abundance for comparison of taxonomic composition across samples. In addition, Spearman correlations were performed to determine relationships between taxonomic and functional profiles and the environmental parameters. Genetic data are available in MG-RAST under the project ID mgp15628. MG-RAST IDs for each sample are described in Supplementary Table 2.

To investigate the complexity of community interactions at each sampling site, we used co-occurrence network analysis. For this, non-random co-occurrence analyses were performed using the Python module ‘SparCC’ (Friedman and Alm, 2012). A table of frequency of hits affiliated to the genus level was used for analysis. For each network, we considered only strong (SparCC > 0.9 or < −0.9) and highly significant (*p* < 0.01) correlations between microbial taxa. The nodes in the reconstructed network represent taxa at the genus level, whereas the edges represent significantly positive or negative correlation between nodes. The analysis of network complexity was based on a set of measures, such as the number of nodes and edges, modularity, the number of communities, average node connectivity, average path length, diameter, and cumulative degree distribution (Newman, 2003). Network visualization and property measurements were calculated with the software Gephi (Bastian et al., 2009).

### Metagenomic assembly and genome reconstruction

We used two different strategies for metagenomic assembly and genomic binning of the eighteen metagenomic datasets from Deception Island volcano. First, reads were assembled using IDBA-ud (Peng et al., 2012) (-mink 50, -maxk 92, -tep 4, -min_contig 1000) and then genomic binning was performed through MaxBin 2.0 (Wu et al., 2016). Contigs were annotated using the Integrated Microbial Genomes & Microbiomes (IMG/M) system (Markowitz et al., 2009) and archived on the JGI/IMG server under Project ID Gs0141992. IMG accession numbers for each sample are described in Supplementary Table 2.

Furthermore, reads were co-assembled using MEGAHIT v. 1.0.2. (Li et al., 2015), discarding contigs smaller than 1000 bp. Then contigs were binned using anvi’o v. 5 following the workflow described by Eren et al. (2015). Reads for each metagenome were mapped to the co-assembly using bowtie2 with default parameters (Langmead and Salzberg, 2012). A contig database was generated using the ‘anvi-gen-contigs-database’. Prodigal (Hyatt et al., 2010) was used to predict open reading frames (ORFs). Single-copy bacterial and archaeal genes were identified using HMMER v. 3.1b2 (Finn et al., 2011). The program ‘anvi-run-ncbi-cogs’ was used to annotate genes with functions by searching for them against the December 2014 release of the Clusters of Orthologous Groups (COGs) database (Galperin et al., 2015) using blastp v2.10.0+ (Altschul et al., 1990). Predicted protein sequences were functionally and taxonomically annotated against KEGG with GhostKOALA (genus_prokaryotes) (Kanehisa et al., 2016). Individual BAM files were profiled using the program ‘anvi-profile’ with a minimum contig length of 4 kbp. Genome binning was performed using CONCOCT (Alneberg et al., 2013) through the ‘anvi-merge’ program with default parameters. We used ‘anvi-interactive’ to visualize the merged data and identify genome bins. Bins were then manually refined using ‘anvi-refine’, and completeness and contamination were estimated using ‘anvi-summarize’.

Bins generated by the assembly and co-assembly approaches were quality checked through CheckM v. 1.0.7 (Parks et al., 2015), which is based on the representation of lineage-specific marker gene sets. Bins were taxonomically classified based on genome phylogeny using GTDB-Tk (Chaumeil et al., 2020).

### Taxonomic and functional annotation of metagenome-assembled genomes (MAGs)

Bins were defined as a high-quality draft (>90% complete, <5% contamination), medium-quality draft (>50% complete, <10% contamination) or low-quality draft (<50% complete, <10% contamination) metagenome assembled-genome (MAG), according to genome quality standards suggested by Bowers et al. (2017). We selected 11 MAGs based on their medium or high-quality and taxonomy, preferably selecting groups related to extremophiles or associated to sulfur and nitrogen metabolisms. Annotation of all predicted ORFs in MAGs was performed using prokka v.14.5 (Seemann, 2014). Further, proteins were compared to sequences in the KEGG Database through GhostKOALA (genus_prokaryotes) (Kanehisa et al., 2016) and in the SEED Subsystem through RASTtk (Brettin et al., 2015). Phenotypes were predicted using the PICA framework (Feldbauer et al., 2015) and PhenDB (https://phendb.csb.univie.ac.at/).

## Results

To investigate links between metabolic potential and genes associated with survival strategies across extreme temperature and geochemical gradients of the Deception Island volcano, we analyzed the metagenomes of a total of eighteen samples, comprising fumaroles with temperatures of 98 °C, 80 °C, 50 °C, and 10 °C, and glaciers with temperatures around 0 °C. Shotgun sequencing of community genomic DNA on 3 lanes of Illumina HiSeq2000 produced a total of 567,410,264 paired-end reads, within which 475,895,996 were filtered by quality (Q>30) for further analyses. A total of 162,755,88 reads were taxonomically annotated as Bacteria, 3,680,020 as Archaea, 2,094,916 as Eukarya and 79,111 as viruses (Supplementary Table 2). The total number of proteins predicted in reads were 296,818,692 (62.3%). Relative abundances of the detected genes were used to compare the metabolic potential and genes related to survival strategies under environmental extremes among fumaroles and glaciers samples.

*De novo* assemblies of the quality-filtered reads generated a total of 543,945 contigs. The prediction of ORFs resulted in 1,396,820 putative genes, 353,731 assigned within Bacteria, 12,034 within Archaea, and 1,557 and 1,534 within Eukarya and viruses, respectively. We used different databases for assembly annotation through JGI/IMG that resulted in 487 putative 16S rRNA genes, 842,798 genes based on the COG database and 304,173 genes based on KEGG (Supplementary Table 2).

### Taxonomic profile of microbial communities on the Deception Island volcano

Through the annotation of reads, we observed that the taxonomic composition in the 98 °C fumarole was distinct in comparison with other fumaroles and glaciers. Archaea were dominant in samples from the 98 °C fumarole (relative abundance between 31.5 and 87.3%), with the most abundant archaeal phyla classified as Crenarchaeota (23.8-79.3%), followed by Euryarchaeota (2.5-7.5%) and Korarchaeota (0.1-0.4%). Firmicutes (3.1-22.4%), Bacteroidetes (0.6-15.3%), Aquificae (0.3-4.6%), and Thermotogae (0.3-1.0%) were also detected in minor proportions in the 98 °C fumarole. Looking at the class level, Thermoprotei, Thermococci, Methanococci, Archaeoglobi, Methanobacteria, Methanopyri, and Methanomicrobia represented the most abundant archaeal classes (>0.1%) in the 98 °C site, and Bacilli, Gammaproteobacteria, Betaproteobacteria, Fusobacteria, Flavobacteria, Aquificae (order Aquificales) and Thermotogae (order Thermotogales) were the dominant classes within Bacteria (Figure 2a).

**Figure 2.**
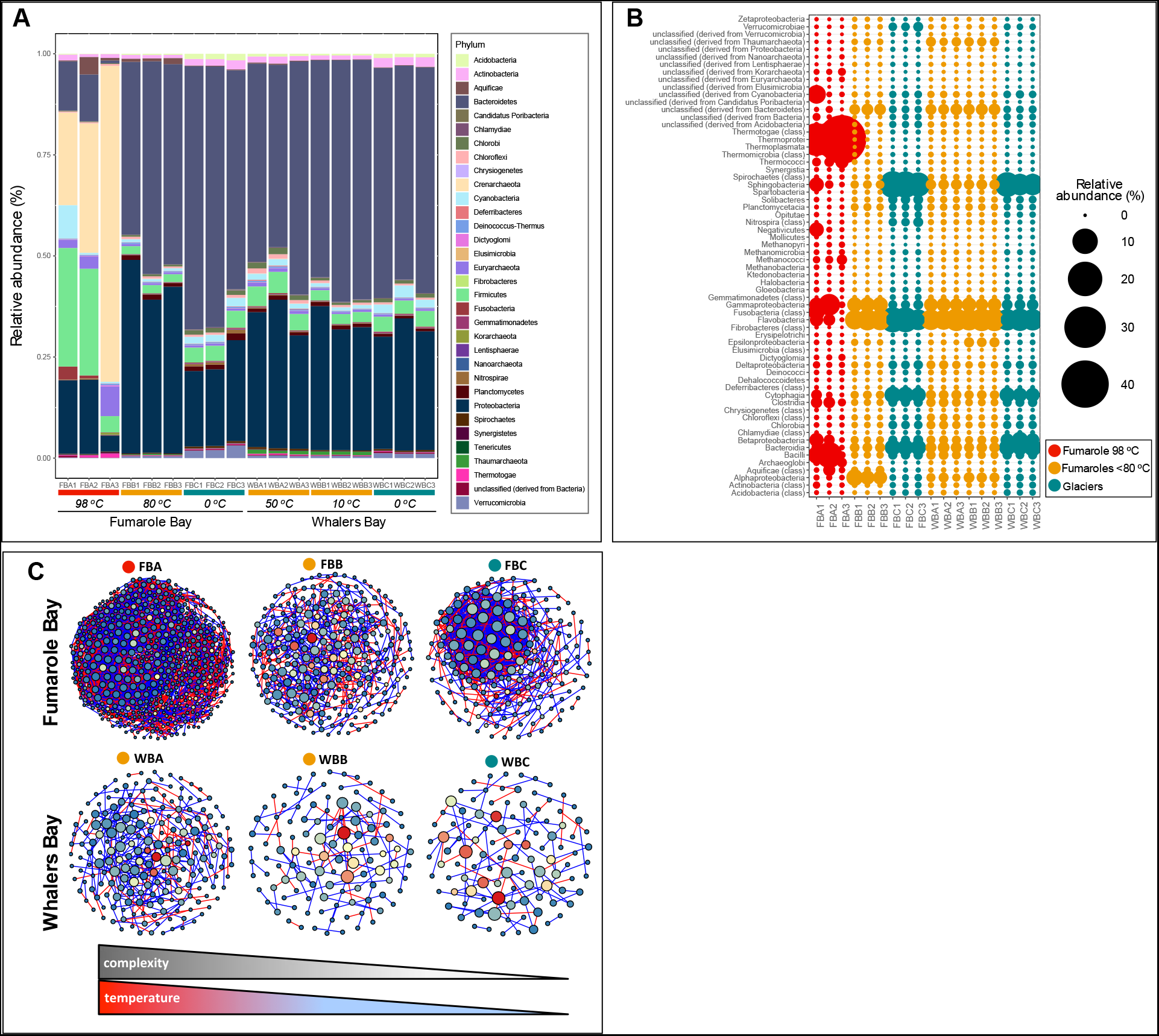
Relative abundances of microbial community taxonomy based on annotation of reads from shotgun metagenomics, represented at phylum (A) and class (B) levels. Environmental temperatures and geothermal sites of each sample are represented. Taxonomy assignments were performed based on best hit classifications and M5NR (non-redundant protein database), with an e-value of <1×10^−5^, minimum 50 bp alignment, and 60% identity. Co-occurrence network analysis at the genus level are represented in (C), grouping triplicates of each sampling point and highlighting the increases of environmental temperatures and complexity. Complexity was calculated based on a set of measures, such as the number of nodes and edges, modularity, the number of communities, average node connectivity, average path length, diameter, and cumulative degree distribution.

Archaea were less dominant in the other samples, with a relative abundance of 0.7-2% in fumaroles <80 °C and 0.4-0.6% in glaciers. Although some dominant phyla were common between <80 °C fumaroles and glaciers (e.g. Bacteroidetes, Proteobacteria, and Firmicutes), less dominant phyla were uniquely distributed according to temperature. For example, Thaumarchaeota was predominantly found in <80 °C fumaroles (0.8-1% for Whalers Bay and 0.2-0.3% in Fumarole Bay). Verrucomicrobia and Acidobacteria were only detected in glaciers (1.2-3.1% and 1-1.6%, respectively) (Figure 2a). The main classes affiliated within the Bacteroidetes phylum were Cytophagia, Flavobacteria and Sphingobacteria, whereas Gamma- and Alphaproteobacteria were the most represented classes within Proteobacteria, followed by Beta-, Delta- and Epsilonbacteria (Figure 2b). Solibacteres was the abundant class within Acidobacteria, and Verrucumicrobiaea within Verrucomicrobia. Thaumarchaeota assignments were not classified at the class level using reads annotation in MG-RAST. The taxonomic annotation of contigs through the IMG/M system showed similar patterns when compared to reads annotation (Supplementary Figure 1).

We then used co-occurrence network analysis to explore the complexity of interactions within the microbial communities in each treatment (Figure 2c). For this, we calculated SparCC correlations between microbial taxa at the genus level based on metagenome reads annotated in MG-RAST. In general, the complexity of the community increased with the temperature. We also noted that communities of Fumarole Bay were more complex than Whalers Bay. The FBA (98 °C) site showed the highest level of complexity and a modular structure, whereas the WBC (0 °C) site had the least complex network. Interestingly, the proportion of positive/negative correlations also changed according to the temperature; at higher temperatures, the proportion is even, while in lower temperatures there was an increase in the number of positive correlations.

### Comparative functional profile of microbial communities on Deception Island volcano

Functional profiles of metagenomes were compared using a multivariate method and hypothesis test, and significant variations in all functional levels were observed between the 98 °C fumarole, <80 °C fumaroles and glaciers. Clear differences between these three distinct sample groups were observed through both the SEED Level 1 profile (Figure 3a) and SEED functional level through PCA ordination (Figure 3b). Further, a distinct pattern between samples from the highest temperature was observed. The prevalent core of functions among Deception samples were “clustering-based subsystems”, “carbohydrates”, “amino acids and derivatives”, “protein metabolism”, “RNA metabolism”, “DNA metabolism” and “cofactors, vitamins, prosthetic groups and pigments”. Significant differences between sample groups and functions from level 1 of SEED, calculated using Fisher’s exact two-sided test and the Newcombe-Wilson method, showed the highest abundance of genes belonging to the categories “DNA metabolism” (*p* = 0.046), “protein metabolism” (*p* = 0.049), and “phages, prophages, transposable elements and plasmids” (*p* = 3.97e-3) in the 98 °C fumarole in comparison with other fumaroles and glaciers. The categories “nitrogen metabolism” (p = 1.07e-4), “photosynthesis” (*p* = 5.07e-3), “sulfur metabolism” (*p* = 0.015), and “metabolism of aromatic compounds” (*p* = 0.036) exhibited the highest significant values in <80 °C fumaroles when compared to the 98 °C fumarole, and “motility and chemotaxis” (*p* = 2.71e-8), “RNA metabolism” (*p* = 0.01), and “protein metabolism” (*p* = 0.023) when compared to glaciers. In contrast, observations for glaciers showed more genes associated with “carbohydrates” and “virulence, disease and defense” categories in comparison with the 98 °C fumarole (*p* = 0.049 and *p* = 0.036, respectively) and <80 °C fumaroles (*p* = 1.68e-6 and *p* = 0.012, respectively) (Figure 3c).

**Figure 3.**
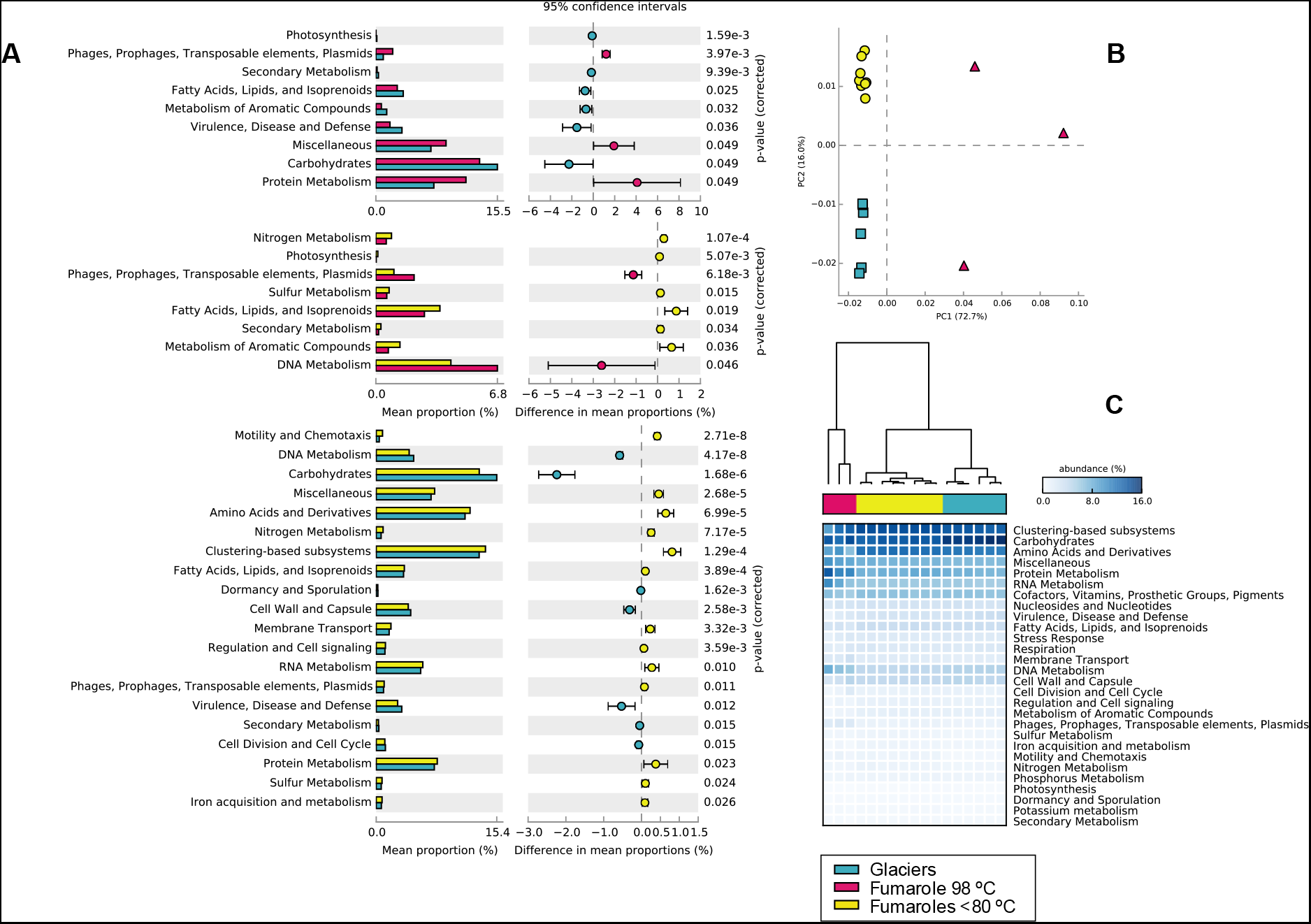
Extended error plots for functional general profiles of microbial communities generated through annotation of metagenomic reads, visualized through STAMP based on SEED subsystems, are represented in (A). The *p* values were calculated using Fisher’s exact two-sided test and the confidence intervals were calculated by the method from Newcombe-Wilson. Differences were considered significant at *p*< 0.05. PCA ordination was performed based on functions at level 3 of the SEED subsystem (B). Heatmap is representing relative abundances of level 1 functions (C). Samples are clustered and colored according to environmental temperature, following the three different groups: 98 °C fumarole, <80 °C fumaroles and glaciers.

### Patterns of metabolic partitioning among extreme temperatures

We observed different partitioning patterns of metabolic diversity according to environmental temperatures (Figure 4). The fumarole with the highest temperature (98 °C) exhibited metabolic potential significantly higher for functions associated with sulfate reduction (*p* < 0.001), dissimilatory nitrite reduction (*p* < 0.001) and carbon dioxide fixation (*p* < 0.001) when compared to other fumaroles and glaciers. Although sulfur metabolism was abundant among all temperatures, different metabolic pathways related to sulfur were observed according to the temperature. While sulfate reduction was prevalent in the highest temperature fumarole, a high number of genes related to inorganic sulfur assimilation (*p* < 0.001) and sulfur oxidation (*p* value was not significant, *p* < 0.1) were detected in <80 °C fumaroles. In general, nitrogen metabolism was dominant in <80 °C fumaroles when compared to other samples, with nitrate and nitrite ammonification (*p* < 0.02), denitrification (*p* < 0.01), nitrogen fixation (*p* < 0.05) and ammonia assimilation (*p* < 0.001) as the prevalent metabolic nitrogen pathways. All fumaroles showed a similar abundance of genes belonging to sulfur oxidation, nitrate and nitrite ammonification, and dissimilatory nitrite reduction. The genetic potential for carbon fixation was much higher in the 98 °C fumarole (*p* < 0.01), whereas photosynthesis was mainly detected in the <80 °C fumaroles and glaciers. In glaciers, the genes identified within carbon metabolism were mainly associated with heterotrophy and central carbon pathways, such as the pentose phosphate pathway and glycolysis, as were respiration and fermentation. The function of carbon storage regulators was significantly higher in <80 °C fumaroles, in addition to the observation of other carbon-related processes, such as photosynthesis, fermentation, and carbon fixation (Figure 4).

**Figure 4.**
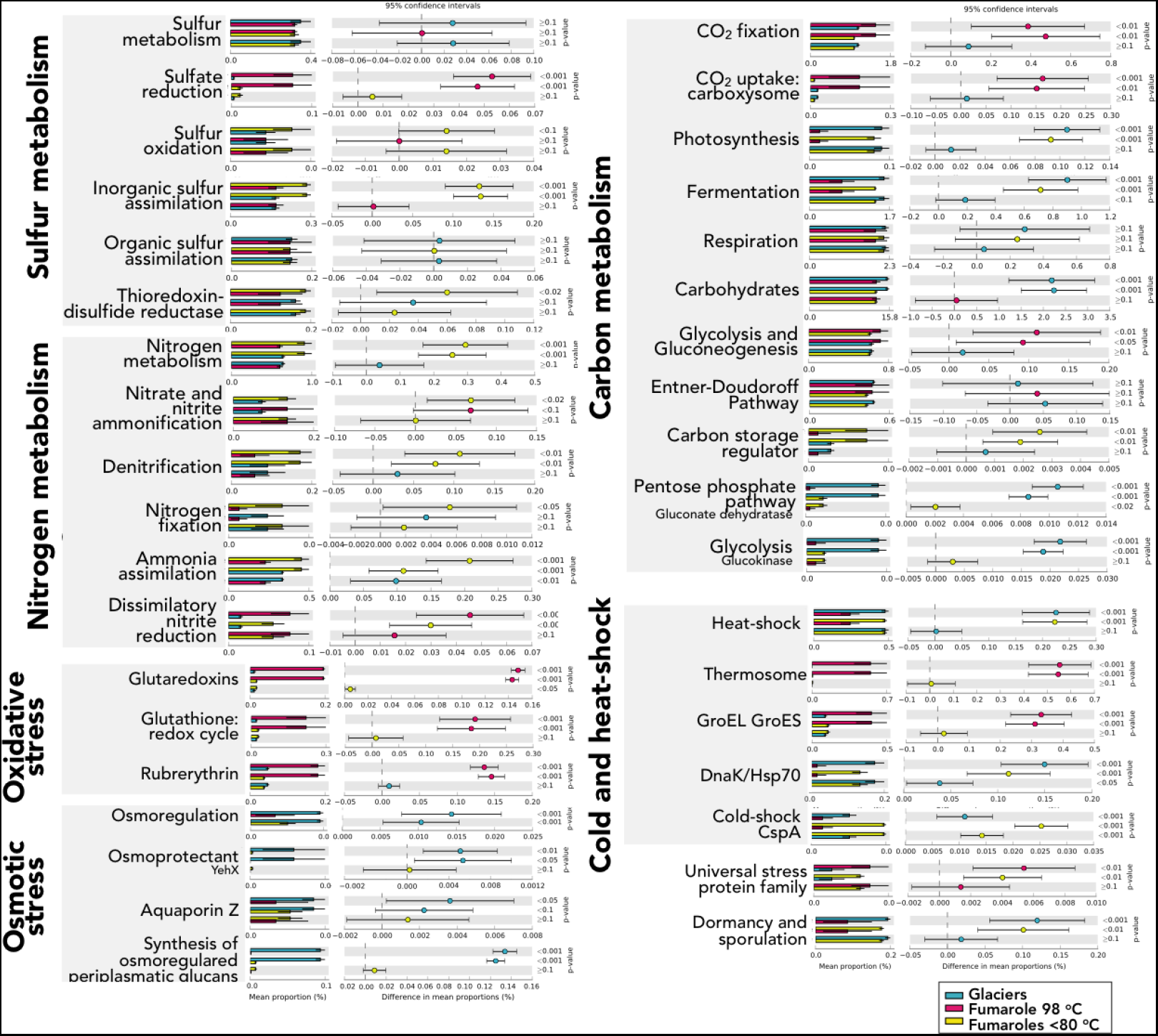
Extended error plots for functional profiles regarding metabolic pathways, including sulfur, nitrogen and carbon metabolisms, and stress response, including oxidative and osmotic, and heat/cold shock responses. Profiles were visualized through STAMP based on annotation of metagenomic reads using SEED subsystems. The *p* values are represented and were calculated using Fisher’s exact two-sided test, with the confidence intervals calculated by the method from Newcombe-Wilson. Samples are clustered and colored according to environmental temperature, following the three different groups: 98 °C fumaroles, <80 °C fumaroles and glaciers.

### Community survival strategies under environmental extremes

To understand community survival strategies under extreme temperature and geochemical gradients, we selected genes in our metagenomes that are involved with stress response, DNA repair, protein biosynthesis, and transport and chemotaxis. Although communities from all samples were equally abundant in genes related to stress response, very distinct patterns of specific responses were observed accordingly environmental temperature (Figure 4). The oxidative stress response was markedly higher in the fumarole at 98 °C, mainly represented by glutaredoxins, glutathione (redox cycle) and rubrerythrin functions (all with *p* < 0.001 in comparison with other samples). Contrastingly, osmotic stress genes were prevalent in glaciers samples, represented mainly by functions such as osmoregulation (*p* < 0.001), osmoprotectant (*yehX*) (*p* = 0.01), aquaporin Z (*p* = 0.05) and synthesis of osmoregulated periplasmatic glucans (*p* < 0.001). The abundance of genes related to heat and cold responses (thermal response) was distinctly distributed among fumaroles and glaciers. General function of heat-shock proteins (including *hsp70*/*dnaK*) were prevalent in glaciers and <80 °C fumaroles (*p* < 0.001), whereas specific archaeal thermal responses dominated the 98 °C fumarole, such as thermosome chaperonin (*p* < 0.001, 0.7% of total relative abundance), as were bacterial and archaeal heat-shocks *groEL*/*groES* (*p* < 0.001). The relative abundance of cold shock *cspA* was higher in <80 °C fumaroles, followed by glaciers (*p* < 0.001). Glaciers and <80 °C fumaroles exhibited the highest abundance of dormancy and sporulation function (*p* < 0.01) and all fumaroles had a prevalence of the universal stress protein family (*p* < 0.01) (Figure 4).

Differences in abundance patterns of DNA repair, protein biosynthesis, transport and chemotaxis were also observed across environmental temperatures (Supplementary Figure 2). Base excision repair, recombination through *recU* and reverse gyrase (all with *p* < 0.01) were the main strategies of DNA positive supercoiling and repair notably found in communities of the highest temperature fumarole (98 °C). Strategies of DNA repair using *uvrABC* complex, recombination through *recA* and photolyase were dominant in <80 °C fumaroles and glaciers (all with *p* < 0.001). Protein biosynthesis genes were dominant in the highest temperature fumarole (98 °C); functions such as universal GTPases (*p* < 0.01) and translation elongation factors in Archaea (*p* < 0.001) were significantly higher when compared to the other samples. Chemotaxis genes were also prevalent in the highest temperature fumarole (98 °C) (*p* < 0.001), as were several transport systems (transport of Ni, Co, and Zn) and ABC transporters (e.g. branched-chain amino acid, oligopeptide and tungstate) (all with at least *p* < 0.05). Mn transport and the ABC transporters of iron and peptides were significantly higher in <80 °C fumaroles (all with at least *p* < 0.05) (Supplementary Figure 2).

### Physical-chemical influence on taxonomic and functional diversity

To identify key environmental drivers of community taxonomy (at phylum level) and function (SEED level 1), Spearman correlations were calculated; then only significant (*p* < 0.05) and strong correlations (*r* > −0.6 or 0.6) were considered. In general, the phyla that positively correlated with temperature were Euryarchaeota, Crenarchaeota, Korarchaeota, Nanoarchaeota, Thermotogae, and Aquificae, whereas several phyla were negatively correlated with temperature (e.g. Acidobacteria, Bacteroidetes, Spirochaetes, Actinobacteria, Verrucumicrobia, Nitrospirae, Deinococcus-Thermus, and Gemmatimonadetes, among others) (Figure 5a). The phyla that negatively correlated with ammonia were Proteobacteria, Thaumarchaeota, Euryarchaeota, and Deferribacteres; those positively correlated were Firmicutes, Acidobacteria, Cyanobacteria, and Spirochaetes, among others. All significant nitrate correlations were positive, including phyla such as Firmicutes, Nitrospirae, Thermotogae, etc. Sulfate showed significant positive correlations with Proteobacteria, Thaumarchaeota, Euryarchaeota, Deferribacteres and Crenarchaeota, and negative correlations with phyla such as Acidobacteria, Cyanobacteria, Actinobacteria, and Verrucomicrobia, among others. Other parameters exhibited positive and negative correlations with several phyla, such as organic matter, organic carbon, B, Cu (uniquely negative correlations), Fe (uniquely negative correlations), Na, K, Ca, Mg and Al (Figure 5a, Supplementary Table 3).

**Figure 5.**
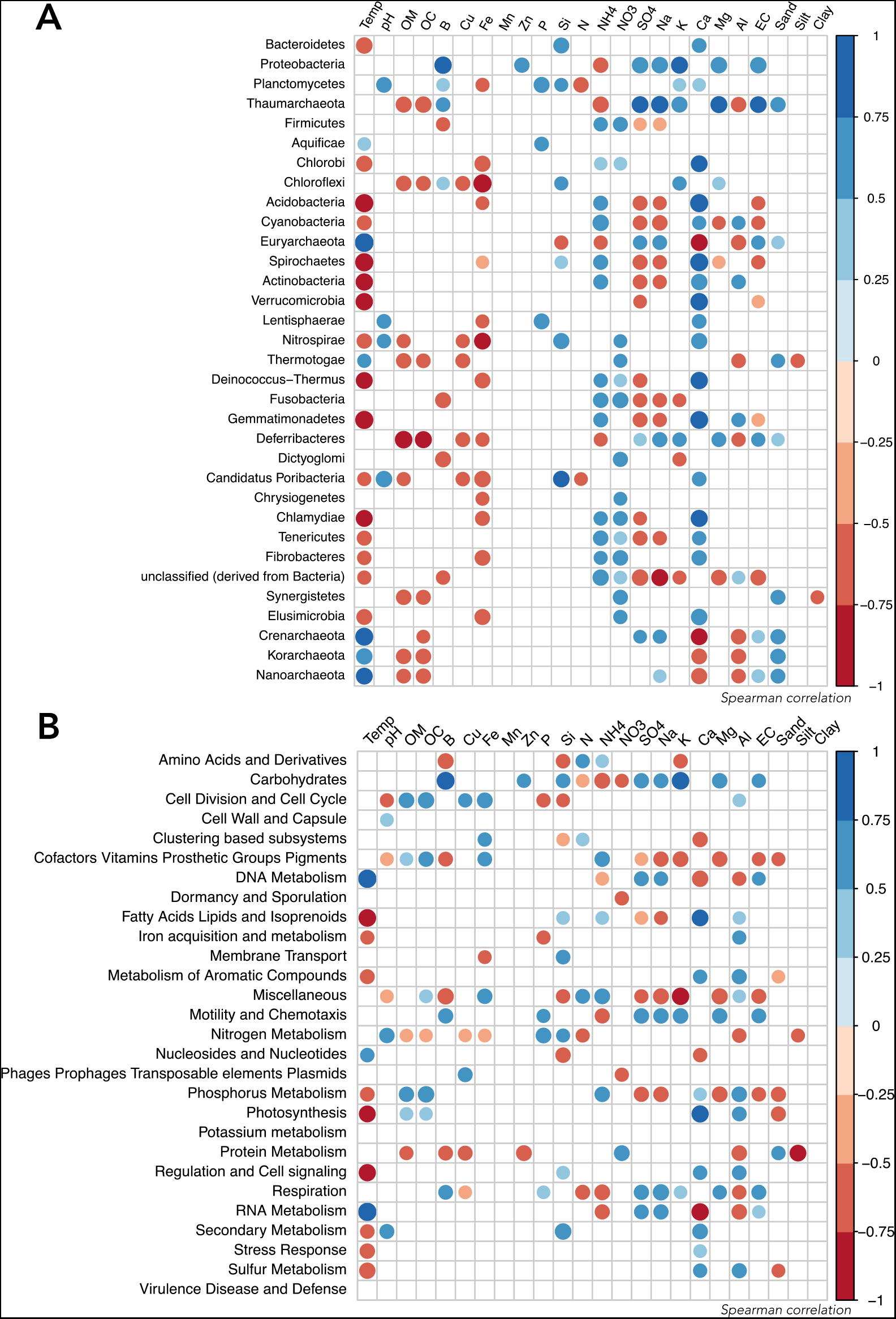
Spearman correlation between taxonomic profile (A) (phylum level) and functional profile (level 1 SEED subsystem) (B) and environmental parameters. Only parameters that exhibited p < 0.05 in a correlation analysis are represented. The environmental parameters are: Temp (temperature), pH, EC (electrical conductivity), B, Cu, Fe, Zn, OM (organic matter), OC (organic carbon), P, Si, Na, K, Ca, Mg, sulfate, nitrogen, ammonia, nitrate, sand, silt, and clay.

The functional categories (SEED level 1) which presented positive correlations with temperature were “DNA metabolism”, “nucleosides and nucleotides” and “RNA metabolism”, and those which exhibited negative correlations were “fatty acids, lipids and isoprenoids”, “iron acquisition”, “metabolism of aromatic compounds”, “phosphorus metabolism”, “photosynthesis”, “secondary metabolism”, “stress response”, and “sulfur metabolism” (Figure 5b). Ammonia was negatively correlated with functions such as “carbohydrates”, “motility and chemotaxis”, “respiration” and “RNA metabolism”, whereas positive correlations comprised functions as “amino acids and derivatives” and “cofactors, vitamins, prosthetic groups and pigments”. Nitrate also presented negative correlations with “carbohydrates”, “dormancy and sporulation” and “phages, prophages, transposable elements and plasmids”. In contrast, sulfate was positively correlated with “carbohydrates”, “DNA metabolism”, “motility and chemotaxis”, “respiration” and “RNA metabolism”. Other parameters exhibited positive and negative correlations with several functions, such as organic matter, organic carbon, B, Cu, Fe, Si, Na, K, Ca, Mg and Al (Figure 5b, Supplementary Table 3).

### Metabolic potential and survival strategies in MAGs

In general, the anvi’o pipeline using co-assembly showed the best binning results for our eighteen metagenomes, generating a total of 158 MAGs. We included in our analyses only 1 MAG produced through the idba-ud assembler and MaxBin binning since this MAG belonged to a taxon (Calditrichia) which was not achieved through the anvi’o pipeline (Supplementary Figure 3, Supplementary Table 4). From the 159 MAGs, 12 were assigned as Archaea through GTDB-Tk and GhostKoala, belonging to Nitrososphaerales (*Candidatus Nitrosocaldus* according with GhostKoala taxonomy) (2), *Nitrosoarchaeum* (1), *Nitrosotenius* (1), *Nitrospumilus* (1), Desulfurococcales (*Aeropyrum* according with GhostKoala taxonomy) (1), Acidilobaceae (1), Pyrodictiaceae (2) and Woesearchaeia (Nanoarchaeota) (3). The bacterial MAGs were classified through GTDB-Tk and GhostKoala as the following phyla: Acidobacteriota (1), Aquificota (2), Bacteroidota (92), Calditrichota (5), Campylobacterota (1), Chloroflexota (3), Cyanobacteriota (1), Firmicutes (1), Nitrospirota (2), Patescibacteria (4) and Proteobacteria (35) (Supplementary Figure 4, Supplementary Table 4). A total of 13 MAGs were considered as high quality and 82 as medium quality drafts.

The MAGs were selected for functional annotation by their quality and based on groups related to extremophiles and associated to sulfur and nitrogen metabolisms. These 11 selected MAGs were assigned as DI_MAG_00003 (*Sulfurimonas*), DI_MAG_00004 (Hydrogenothermaceae/*Persephonella*), DI_MAG_00006 (Promineofilaceae/*Candidatus Promineofilum*), DI_MAG_00010 (Caldilineaceae/*Caldilinea*), DI_MAG_00011 (Thermonemataceae), DI_MAG_00019 (Chitinophagaceae), DI_MAG_00020 (Pyrodictiaceae/*Pyrodictium*), DI_MAG_00021 (Dojkabacteria), DI_MAG_00022 (Woesearchaeia/archaeon GW2011_AR20), DI_MAG_00049 (Nitrososphaerales/*Candidatus Nitrosocaldus*) and DI_MAG_FBB2_12 (Calditrichia) (Table 1).

**Table 1.** List of the 11 selected MAGs and their taxonomic classification based on GTDB-Tk and GhostKoala. Characteristics of total genome length, N50, GC content, redundancy and completeness (based on CheckM), and the genome quality status, are described.

| MAG Code | KEGG/Ghost<br>Koala taxonomy | GTDB-Tk taxonomy | Total<br>Length | Contigs<br>number | N50 | GC<br>content | Completeness | Redundancy | Draft<br>Quality |
| --- | --- | --- | --- | --- | --- | --- | --- | --- | --- |
| DI_MAG_00003 | *Sulfurimonas* | Bacteria; Campylobacterota;<br>Campylobacteria;<br>Campylobacterales; Thiovulaceae;<br>*Sulfurimonas* | 1,912,170 | 94 | 32,276 | 39.42 | 97.95 | 0.41 | High |
| DI_MAG_00004 | *Persephonella* | Bacteria; Aquificota; Aquificae;<br>Hydrogenothermales;<br>Hydrogenothermaceae | 1,682,998 | 67 | 44,923 | 30.32 | 98.37 | 0.22 | High |
| DI_MAG_00006 | *Candidatus*<br>*Promineofilum* | Bacteria; Chloroflexota;<br>Anaerolineae; Promineofilales;<br>Promineofilaceae | 4,660,983 | 56 | 102,288 | 52.27 | 92.73 | 2.00 | High |
| DI_MAG_00010 | *Caldilinea* | Bacteria; Chloroflexota;<br>Anaerolineae; Caldilineales;<br>Caldilineaceae | 4,473,267 | 92 | 90,197 | 57.82 | 96.36 | 0.91 | High |
| DI_MAG_00011 | Unknown | Bacteria; Bacteroidota; Bacteroidia;<br>Cytophagales; Thermonemataceae | 2,518,210 | 152 | 22,003 | 49.00 | 88.43 | 0.55 | High |
| DI_MAG_00019 | Unknown | Bacteria; Bacteroidota; Bacteroidia;<br>Chitinophagales; Chitinophagaceae | 3,082,789 | 324 | 10,795 | 37.53 | 62.85 | 7.88 | Medium |
| DI_MAG_00020 | *Pyrodictium* | Archaea; Crenarchaeota;<br>Thermoprotei; Desulfurococcales;<br>Pyrodictiaceae | 1,071,537 | 28 | 68,816 | 47.76 | 75.27 | 0.47 | Medium |
| DI_MAG_00021 | Unknown | Bacteria; Patescibacteria;<br>Dojkabacteria | 734,971 | 8 | 176,471 | 31.53 | 77.27 | 1.72 | Medium |
| DI_MAG_00022 | archaeon<br>GW2011_AR20 | Archaea; Nanoarchaeota;<br>Woesearchaeia | 1,155,662 | 40 | 83,443 | 44.63 | 79.44 | 0.00 | Medium |
| DI_MAG_00049 | *Candidatus*<br>*Nitrosocaldus* | Archaea; Crenarchaeota;<br>Nitrososphaeria; Nitrososphaerales | 982,006 | 145 | 6,877 | 32.09 | 60.52 | 1.94 | Medium |
| DI_MAG_FBB2_12 | *Caldithrix* | Bacteria; Calditrichota; Calditrichia | 2,931,548 | 176 | 21,244 | 39.10 | 95.54 | 0.00 | High |

We identified in the high-quality DI_MAG_00004 (Hydrogenothermaceae/*Persephonella*, ~ 97% completeness) genes for nitrate reduction (*narGHI* and *nirA*), denitrification (*narGHI*), nitrification (*narGH*), sulfate reduction (*sat*, *cysH*, *sir*), sulfur and thiosulfate oxidation (*soxAXBYZ*), and incomplete pathways for carbon fixation (Figure 6a). This MAG had several genes associated with stress response, especially oxidative stress (e.g. superoxide reductase and dismutase, rubrerythrin and rubredoxin) and thermal response (e.g. *groES*, *hsp20* and *hspR*), as different DNA repair mechanisms, including photolyase repair (Figure 6b). In general, genes involved with the nitrogen cycle were identified in almost all selected MAGs, except for MAGs DI_MAG_00020, DI_MAG_00021, and DI_MAG_00022. Sulfate reduction genes were also detected in different selected MAGs, except for MAGs DI_MAG_00020, DI_MAG_00021, DI_MAG_00022 and DI_MAG_FBB2_12. All MAGs had incomplete pathways for carbon fixation, except for DI_MAG_00004 and DI_MAG_00021 (Figure 6a).

**Figure 6.**
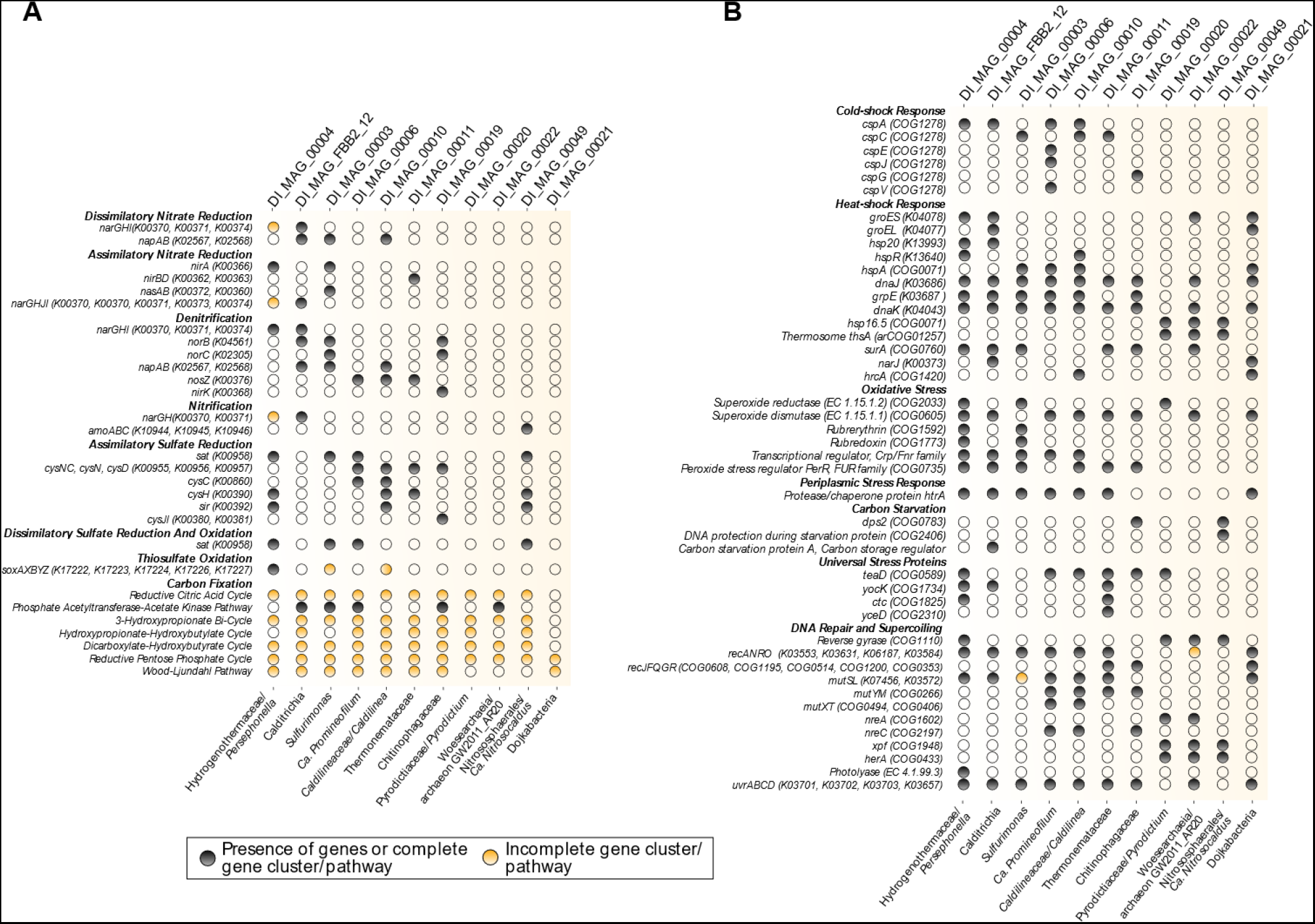
Functional annotation of the 11 selected metagenome-assembled genomes (MAGs), including metabolic potential (A) and adaptive strategies (B). A black circle represents the presence of genes or complete gene cluster/pathway, and a yellow circle represents incomplete gene cluster or pathway. MAGs codes are represented on the upper side of figures, whereas their taxonomic classification based on GTDB-Tk and GhostKoala are at the bottom. Genes are presented here with identifiers of KEGG Orthology (KO), Clusters of Orthologous Groups (COG) or Enzyme Commission numbers (EC).

Different cold-shock genes were detected among MAGs; DI_MAG_00006 was the one which presented more *csp* genes. We did not find any *csp* genes in archaeal MAGs (DI_MAG_00020, DI_MAG_00022, and DI_MAG_00049). However, we observed genes in all selected MAGs that were related to different heat-shock responses, including *groEL/groES* genes in DI_MAG_00004, DI_MAG_00020, and DI_MAG_00021. Thermosome (*thsA*) and reverse gyrase genes were identified in all the selected MAGs assigned as Archaea (DI_MAG_00020, DI_MAG_00022, and DI_MAG_00049). Although all MAGs showed the potential presence of oxidative stress response (except DI_MAG_00049), rubrerythrin and rubredoxin genes were only observed in DI_MAG_00004 and DI_MAG_00003. Different DNA repair mechanisms were identified in selected MAGs, such as several recombination genes (*rec* genes), DNA mismatch repair (*mut* genes), nucleotide excision repair (*uvr* genes), double-strand break repair (*her*A, only in archaeal-selected MAGs) and photolyase repair (only in DI_MAG_00004) (Figure 6b).

## Discussion

The primary goal of our study was to unveil how communities functionally respond to the combination of environmental factors typical of polar marine volcanoes. Our results show that regardless of proximity between fumaroles and glaciers on Deception Island, the community function is strongly driven by the combination of contrasting environmental factors, as occurred similar to what we previously observed for community composition and diversity (Bendia et al., 2018b). We detected some bacterial groups present in both glacier and fumarole sediments (most notably the phyla Proteobacteria, Firmicutes, and Bacteroidetes), despite the strong gradients in temperature, geochemistry and salinity. In addition, we observed specific groups that varied according to the environmental temperature: the hyperthermophilic members belonging to Crenarchaeota/Thermoprotei, Aquificae and Thermotoga phyla in the 98 °C fumarole, Thaumarchaeota in <80 °C fumaroles, and Acidobacteria and Verrucomicrobia in glaciers. These patterns are consistent with previous work carried out on Deception Island using the same sample set for diversity analysis (16S rRNA gene sequencing) (Bendia et al., 2018b), except for the Aquificae and Thermotogae phyla, which were not detected by that method. Furthermore, our taxonomic patterns were also consistent with a previous report that observed similar members along a temperature gradient ranging from 7.5 to 99 °C in geothermal areas in Canada and New Zealand (Sharp et al., 2014).

Surprisingly, our network analysis showed that the community interaction in the hottest fumarole (98 °C) was more complex and presented fewer positive interactions when compared to the lowest temperatures, in contrast to previous studies that showed that community complexity decreases with temperature increase (Cole et al., 2013; Merino et al., 2019; Sharp et al., 2014). Our results suggest that hyperthermophilic temperatures on Deception probably trigger ecological interactions between community members to modulate their resistance and resilience when facing strong environmental stressors. Similar patterns of community interaction have been previously observed in stressful conditions in the Atacama Desert (Mandakovic et al., 2018) and with increasing temperature in anaerobic digestion (Lin et al., 2016), although these environmental conditions are different from those found on Deception Island.

Correlation with environmental drivers varied among both taxonomic and functional groups. For example, groups positively influenced by temperature, sulfate, and sodium were those mainly abundant in fumaroles, while groups and functions prevalent in glaciers were positively correlated with ammonia. These results indicate that the mosaic of environmental parameters shapes both taxonomic and functional diversity of microbial communities. Indeed, we observed a partition of metabolic diversity among the steep environmental gradients on Deception Island. Unlike previous studies carried out at hydrothermal vents which pointed to metabolic functional redundancy at the community level (Galambos et al., 2019; Reveillaud et al., 2016), Deception communities showed metabolic heterogeneity across the sharp temperature gradient. The observation of functional redundancy despite the taxonomic variation has been observed in several environments such as venting fluids from the Mariana back-arc, cold subseafloor ecosystems, freshwater and gut microbiomes (Louca et al., 2016; Trembath-Reichert et al., 2019; Tully et al., 2018; Turnbaugh and Gordon, 2009; Várbíró et al., 2017). The metabolic heterogeneity observed in our results indicates that microbial communities on Deception harbor a remarkably diverse genetic content that reflects the strong selective pressures caused by a remarkable interaction between the volcanic activity, the marine environment, and the cryosphere.

The functional pattern clustered the samples by temperature, rather than by geographic location, and showed that microbial communities on Deception Island are grouped by 98 °C fumarole, <80 °C fumaroles and glaciers. The predominant metabolic potential in the hottest fumarole (98 °C) was mostly associated with reductive pathways, such as sulfate reduction, ammonification, and dissimilatory nitrite reduction, and carbon fixation. We suggest that the hydrogen sulfide emissions and hyperthermophilic conditions of this fumarole (98 °C) (Somoza et al., 2004) may decrease the dissolved oxygen even in the superficial sediment layers, creating a steep redox gradient and preferably selecting microorganisms with reductive and autotrophic pathways. In addition, communities from the hottest fumarole (98 °C) exhibited several genes related to different adaptive strategies, such as those associated with oxidative stress, specific archaeal heat-shock responses, base excision repair, recombination (*recU*), reverse gyrase, protein biosynthesis, chemotaxis, and ABC transporters. This reflects its primaries stress factors, including the fumarolic production of hydrogen sulfide, which has a strong reductive power capable of causing oxidative stress, and hyperthermophilic temperature that induces disturbance to metabolic processes and cell-component denaturation (Hedlund et al., 2015; Merino et al., 2019). Enrichment in genes involved with chemotaxis was also observed in metagenomes from hydrothermal vents at Juan de Fuca Ridge (Xie et al., 2011), but different DNA repair mechanisms were found when compared to Deception metagenomes. Different types of ABC transporters were also detected in Ilheya hydrothermal fields (Wang and Sun, 2017); reverse gyrase and thermosome mechanisms have often been described in several groups of hyper(thermophilic) Archaea (Forterre et al., 2000; Lemmens et al., 2018; Lulchev and Klostermeier, 2014).

In contrast, <80 °C fumaroles were dominated by genes involved with different energetic and chemolithotrophic pathways: sulfur oxidation, ammonification, denitrification, nitrogen fixation, and dissimilatory nitrite reduction. This suggests a trend for both reductive and oxidative pathways, as well as metabolic versatility and complex biogeochemical processes at the local community level. Although genes related to sulfur and nitrogen pathways were detected in glaciers, the majority of potential pathways for glacier communities were related to carbon metabolism and heterotrophy. This lowest metabolic diversity can be explained by the decrease of marine and volcanic geochemicals (e.g. sulfate) towards glaciers (Supplementary Table 1), making these substrates unavailable for exploiting different energy sources, as occurs in fumaroles. The <80 °C fumaroles and glacier communities harbored mechanisms for both heat and cold-shock genes, dormancy and sporulation functions, and DNA repair mechanisms through *uvrABC* complex, *recA*, and photolyase. Diverse survival strategies in <80 °C fumaroles and glaciers might be explained by community exposure to fluctuating temperatures and redox conditions that are more variable when compared to the stability of hottest fumarole, which maintains the hyperthermophilic temperatures and hydrogen sulfide emissions for long periods. Further, glacier communities exhibited more genes associated with osmotic stress, which reflects the low liquid water availability due to the predominant freezing conditions of the Antarctic ecosystems (Wei et al., 2016).

Although several studies have shown a quantitative decrease in microbial diversity as temperature increases in both geothermal and hydrothermal ecosystems (Cole et al., 2013; Sharp et al., 2014), little is known about how temperature affects ecosystem functioning due to inhibition of key metabolic enzymes or pathways (Hedlund et al., 2015). Despite the limitation of metagenomics in revealing the truly active microbial metabolic pathways, our results increase understanding of the potential temperature limits on different microbial metabolism at the community level and encourage more studies to elucidate the direct effect of temperature extremes on specific biogeochemical processes in Antarctic volcanic ecosystems.

The 159 MAGs recovered from the Deception Island volcano comprised a broad phylogenetic range of archaeal and bacterial phyla. The 11 MAGs selected for annotation included hyperthermophilic and thermophilic lineages, as well as lineages containing homologs of different predicted sulfur and nitrogen pathways, and archaeal groups underrepresented in genome data, such as *Ca. Nitrosocaldus* and Nanoarchaeota/Woesearchaeia. Since *Ca. Nitrosocaldus* was previously reported only in terrestrial geothermal environments (Abby et al., 2018; Daebeler et al., 2018; Torre et al., 2008), their presence on Deception fumaroles represents a novel outcome for the ecological distribution of thermophilic ammonia-oxidizing Archaea and encourages further investigation to better understand their role in marine volcanic ecosystems. Furthermore, the majority of our selected MAGs are equipped with gene-encoding proteins that protect cells against several stressful conditions, including cold and heat-shock, carbon starvation, oxidative and periplasmic stress, and DNA damage, likely enabling survival and adaptation of these microorganisms to a broad combination of extreme parameters. One of our MAGs was closely related to archaeon GW2011_AR20, which is an uncultivated and underrepresented Nanoarchaeota/Woesearchaeia member described previously in aquifer samples and appears to have a symbiotic or pathogenic lifestyle due to the small genome size and lack of some biosynthesis pathways (Castelle et al., 2015). The genome analysis of our Woesearchaeia MAG (archaeon GW2011_AR20, DI_MAG_00022) suggests a novel putative thermophilic lifestyle or at least a potential heat tolerance for this lineage due to the (i) lack of cold-shock genes, (these genes are mostly absent in the genomes of thermophilic archaea, while usually present in psychrophilic/mesophilic archaeal members (Cavicchioli, 2006; Giaquinto et al., 2007), and (ii) the presence of reverse gyrase, thermosome and other heat-shock genes (e.g. *groES*) that are essentially related to (hyper)thermophiles and heat response. Although these heat-shock genes were also detected in some mesophilic archaeal lineages within Halobacteria, Thaumarchaeota, and *Methanosarcina* spp. (Lemmens et al., 2018), reverse gyrase is the only protein found ubiquitously in hyperthermophilic organisms, but absent in mesophiles (Catchpole and Forterre, 2019), pointing to this Woesearchaeia member as a likely thermophile or hyperthermophile.

## Conclusion

By combining the annotation of reads and contigs together with genome reconstruction from metagenomic data, we provide the first genetic and genomic evidence that microorganisms inhabiting the Deception Island volcano possess a variety of adaptive strategies and metabolic processes that are shaped by steep environmental gradients. We observed that hyperthermophilic temperatures (98 °C) preferably select microorganisms with reductive and autotrophic pathways, while communities from fumaroles <80 °C show a high metabolic versatility with both reductive and oxidative pathways, and glaciers harbor communities with metabolic processes especially related to carbon metabolism and heterotrophy. Survival strategies of microorganisms from the hottest fumarole are very specialized in responding to the hyperthermophilic temperatures and oxidative stress, while <80 °C fumaroles and glacier communities possesses a variety of strategies that are capable of responding to fluctuating redox and temperature conditions. We found more complex and negative interactions among the communities from the hottest fumarole (98 °C), which indicate that the strong environmental stressors probably trigger competitive associations among community members. Furthermore, through the reconstruction of MAGs, we were able to clarify a putative novel thermophilic lifestyle for a Woesearchaeia member and a marine lifestyle for a *Ca. Nitrosocaldus* lineage. Our work represents, as far as we know, the first study to reveal through shotgun metagenomics the response of microbial functional diversity to the extreme temperature gradient (0 to 98°C) of an Antarctic volcano. Furthermore, our study was one of the first to recover MAGs from these ecosystems and it provides new insights regarding the metabolic and survival capabilities of different extremophiles inhabiting the Antarctic volcanoes.

## Author Contributions

AB collected the samples, conceived and designed the experiments, performed the experiments, analyzed the data, performed bioinformatic analysis, wrote the paper, and prepared figures and tables. LL contributed to discussion of metagenome-assembled genome analysis, discussed the data, wrote the paper, and reviewed drafts of the paper. LM performed network analysis, wrote the paper, and reviewed drafts of the paper. CS discussed the data, wrote the paper, and reviewed drafts of the paper. BB discussed the data, wrote and reviewed drafts of the paper. VP conceived and designed the experiments, contributed reagents, materials, and analysis tools, and wrote and reviewed drafts of the paper.

## Acknowledgments

We thank the captain and the crew of the research polar vessel Almirante Maximiano, Dr. Luiz Henrique Rosa, Dr. Wânia Duleba, and Dr. Antônio Carlos Rocha Campos for their support in sampling. We are very thankful to LECOM’s research team and Rosa C. Gamba for their scientific support.

## Supplementary Figures and Tables

Figure S1. Relative abundances of microbial community taxonomy based on annotation of contigs, represented at the phylum level. Contigs were constructed through IDBA-ud and annotated using the Integrated Microbial Genomes & Microbiomes (IMG/M) system.

Figure S2. Extended error plots for functional profiles regarding DNA repair, helicase and topoisomerase, protein biosynthesis, and transport and chemotaxis. Profiles were visualized through STAMP based on annotation of metagenomic reads using SEED subsystems. The *p* values are represented and were calculated using Fisher’s exact two-sided test, with the confidence intervals calculated by the method from Newcombe-Wilson. Samples are clustered and colored according to environmental temperature, following the three different groups: 98 °C fumarole, <80 °C fumaroles and glaciers.

Figure S3. A circular view of the 158 metagenome-assembled genomes (MAGs) that were recovered through anvi’o v. 5 pipeline and are represented with the mean coverage of contigs, and the MAGs redundancy, completeness, GC content, total reads mapped and number of SNVs reported. The clustering dendrogram in the center shows the hierarchical clustering of contigs based on their sequence composition, and their distribution across samples.

Figure S4. Heatmap representing the 13 high quality and 82 medium quality MAGs based on read mapping per sample with Z-score. Samples are clustered and colored according to environmental temperature, following the three different groups: 98 °C fumarole, <80 °C fumaroles and glaciers.

Table S1. Physical-chemical parameters data per sample including temperature, pH, EC (electrical conductivity), B, Cu, Fe, Zn, OM (organic matter), OC (organic carbon), P, Si, Na, K, Ca, Mg, sulfate, nitrogen, ammonia, nitrate, sand, silt, and clay.

Table S2. General information about reads and contigs annotation.

Table S3. *P*-values of Spearman correlations comparing taxonomic and functional profiles with the environmental data.

Table S4. A complete list of all reconstructed MAGs with their characteristics: taxonomic classification based on GTDB-Tk and GhostKoala, the total genome length, N50, GC content, redundancy and completeness, based on anvi’o and CheckM, and the genome quality status.

